# Antifungal and Antibiofilm Activity of a Thermostable Lipid Transfer Protein-Enriched Fraction from *Capsicum annuum* L. Seeds against *Candidozyma auris*

**DOI:** 10.64898/2026.11.24.768040

**Authors:** Sofía Macarena Vera González, Constanza Larrondo, Santiago Claver, Jordi Moreno-Romero, Javier Garcia-Pardo, W. David Obregón

**Author notes:** Correspondence (J.G.-P.) and; (J.G.-P.); (ext. 57) (W.D.O.).

## Abstract

The emergence of multidrug-resistant *Candidozyma auris* (*C. auris*) has intensified the search for antifungal molecules with alternative structures and mechanisms of action. Here, we investigated *Capsicum annuum* L. seeds as a source of thermostable antifungal peptides targeting this pathogen. Thermal fractionation of a crude aqueous seed extract (RBPCE) yielded a heat-treated fraction (RBPHT) enriched in low-molecular-weight proteins. LC– MS/MS identified the type I and type II non-specific lipid-transfer proteins Ca-nsLTP1 and Ca-nsLTP2 as its predominant constituents. Compared with RBPCE, RBPHT displayed increased trypsin-inhibitory potency. RBPHT inhibited *C. albicans*, *C. krusei*, and *C. auris* isolates from clades I, III, and IV, including isolates showing no detectable response to fluconazole under the conditions tested. Against a representative South American clade IV isolate, the nsLTP-enriched fraction exhibited minimum inhibitory and fungicidal concentrations of 180 and 360 µg/mL, respectively. RBPHT also suppressed early biofilm establishment by *C. albicans* and *C. auris* and markedly reduced viable biofilm-associated populations when applied after initial cell adhesion. These findings provide the first evidence of antifungal and antibiofilm activity by a *Capsicum* seed-derived nsLTP-enriched fraction against *C. auris* and position Ca-nsLTP1 and Ca-nsLTP2 as promising plant-derived candidates for further antifungal development.

## 1. Introduction

Over the past decade, *Candidozyma auris* (formerly *Candida auris*; hereafter *C. auris*) has emerged as a multidrug-resistant fungal pathogen responsible for healthcare-associated infections with high mortality worldwide [1–3]. First described in 2009 following its isolation from the ear canal of a patient in Japan [4], *C. auris* has since spread rapidly across healthcare systems on multiple continents and has been reported in more than 50 countries [1,5]. Whole-genome sequencing initially classified circulating isolates into five major genomic clades associated with South Asia (clade I), East Asia (clade II), Africa (clade III), South America (clade IV), and Iran (clade V), each of which appears to have emerged independently [5]. More recently, a sixth genetically distinct clade has been identified among isolates from Singapore and Bangladesh [6]. Across these lineages, clinical manifestations range from superficial infections to invasive candidemia, with reported mortality rates of approximately 30–60% [7–9] and potentially higher rates among vulnerable patient populations [7,10].

The epidemiological success of *C. auris* is associated with several distinctive biological characteristics, including its ability to colonize human skin, tolerate elevated temperatures and salinity, persist for prolonged periods on abiotic surfaces, and withstand commonly used disinfectants [2,11–15]. Its capacity to form biofilms further contributes to environmental persistence and reduced antifungal susceptibility [15]. Biofilm development begins with the adhesion of planktonic yeast cells to a surface, followed by proliferation, extracellular matrix production, maturation, and eventual dispersal [11]. Unlike *Candida albicans* (*C. albicans*), which forms highly filamentous biofilms, mature *C. auris* biofilms predominantly consist of densely packed yeast cells embedded within an extracellular matrix [16]. Cells released from mature biofilms may colonize new anatomical or environmental niches, thereby facilitating persistence and transmission. Moreover, sessile cells within *C. auris* biofilms are generally less susceptible to azoles, echinocandins, and some amphotericin B formulations than their planktonic counterparts [17]. Together, its antifungal resistance, environmental persistence, and capacity to form resilient biofilms highlight the need to identify antifungal agents with alternative chemical scaffolds and mechanisms of action.

Natural products used in traditional medicine have historically provided structurally and functionally diverse molecules for antimicrobial drug discovery [18]. Within this context, the Solanaceae family is particularly relevant because of its longstanding importance in traditional medicine and human nutrition [19–21]. The genus *Capsicum* exemplifies this dual nutritional and medicinal relevance. Its species, commonly known as peppers, have been used in traditional medical practices across diverse geographical and cultural settings to manage a wide range of conditions [22–24]. Native to Central and South America, the genus comprises approximately 43 recognized species, five of which are domesticated: *Capsicum annuum* L., *Capsicum frutescens* L., *Capsicum chinense* Jacq., *Capsicum baccatum* L., and *Capsicum pubescens* Ruiz & Pav [25,26]. Following European contact with the Americas, cultivated peppers were disseminated through Spanish and Portuguese maritime trade networks to Europe, Africa, and Asia, where they were progressively incorporated into local food systems and medicinal traditions [22,27].

The medicinal and antimicrobial properties of *Capsicum* species have been attributed to a diverse repertoire of bioactive molecules, including capsaicinoids, phenolic compounds, and plant antimicrobial peptides (AMPs) distributed across different plant tissues [28–30]. Among these, AMPs are attractive candidates for antifungal development because many combine a low molecular weight (typically <10 kDa) with compact, disulfide-stabilized structures that confer resistance to chemical and proteolytic degradation [31].

Over the past two decades, several classes of AMPs have been isolated from *Capsicum* species, including protease inhibitors [30], defensins [32–34], thionin-like peptides [35], and non-specific lipid transfer proteins (nsLTPs) [36–39]. Among them, nsLTPs constitute a structurally distinctive family of small, generally basic, cysteine-rich proteins belonging to the pathogenesis-related protein 14 family [40]. Typically, their primary sequences contain a conserved eight-cysteine motif (8CM), with the general arrangement C–Xₙ–C–Xₙ–CC–Xₙ– CXC–Xₙ–C–Xₙ–C [41,42]. These cysteine residues form four disulfide bonds that stabilize a compact α-helical scaffold surrounding an internal hydrophobic cavity capable of accommodating diverse lipid ligands [42]. nsLTPs were initially classified as type I and type II proteins, with approximate molecular masses of 9 and 7 kDa, respectively. Subsequent phylogenetic classifications expanded the family to ten types—types I, II, C, D, E, F, G, H, J, and K—based on sequence similarity, cysteine spacing, intron position, and the presence of glycosylphosphatidylinositol-anchoring signals [41,43]. Although originally named for their ability to transfer lipids between membranes *in vitro*, nsLTPs participate in broader physiological processes, including cuticle formation, plant development, responses to abiotic stress, and defence against pathogens [44,45].

The antifungal activity of different plant nsLTPs was initially demonstrated mainly against phytopathogenic fungi, but increasing evidence indicates that some family members are also active against opportunistic human-pathogenic yeasts. In *C. annuum*, an nsLTP-containing seed fraction inhibited the growth of *Candida albicans* and *Candida tropicalis*, altered cell morphology, and impaired glucose-stimulated extracellular acidification [39]. Subsequent purification of the approximately 9-kDa Ca-LTP1 from *C. annuum* seeds confirmed its antifungal activity against *C. tropicalis* and demonstrated permeabilization of the yeast plasma membrane [36]. Additional studies of *C. chinense* seeds have further identified multiple approximately 9-kDa nsLTP isoforms with marked structural stability and the capacity to interact with model lipid membranes [46]. More recently, the nsLTP2 CaCLTP2, isolated from *C. annuum* leaves, inhibited several *Candida* species, including *C. albicans*, *C. tropicalis*, and *C. parapsilosis* [37].

Based on these recent observations, we hypothesized that *C. annuum* seeds may contain bioactive peptides with activity against *C. auris*. We therefore evaluated the antifungal potential of crude and nsLTP-enriched aqueous pepper seed extracts against clinically relevant *C. auris* isolates. Their effects on fungal growth, viability, adhesion, and biofilm development were examined. Our findings establish pepper seeds as an underutilized source of thermostable AMPs with antifungal and antibiofilm activity against the emerging multidrug-resistant pathogen *C. auris*.

## 2. Materials and Methods

### 2.1 Plant material and chemicals

Red bell pepper fruits (*Capsicum annuum* L.; Solanaceae) were obtained from local producers in La Plata, Buenos Aires, Argentina, in January 2025. The seeds were manually separated, washed with distilled water, dried for 24 h at 30 °C and stored at −20 °C until use. Sodium chloride, Coomassie Brilliant Blue G-250, N,N,N′,N′-tetramethylethylenediamine (TEMED), tris(hydroxymethyl)aminomethane (Tris), sodium dodecyl sulfate (SDS), bovine serum albumin (BSA), β-mercaptoethanol, N-benzoyl-DL-arginine-p-nitroanilide (BApNA), 4-nitrophenyl-β-D-glucopyranoside (PNPG), and trypsin, were purchased from Sigma-Aldrich (St. Louis, MO, USA). Commercial antifungal discs containing fluconazole (25 µg/disc), caspofungin (5 µg/disc), nystatin (50 µg/disc), or amphotericin B (10 µg/disc), were obtained from Rosco Diagnostica ApS, Albertslund, Denmark. Unless otherwise indicated, all reagents were of analytical grade.

### 2.2 Microbial strains

The yeast strains used in this study included *C. albicans* and *Candida krusei* (*C. krusei*) from the microbial collection of the Centro de Investigación de Proteínas Vegetales (CIPROVE, Argentina). Clinical isolates of *C. auris* belonging to clades I, III, and IV were provided by the Administración Nacional de Laboratorios e Institutos de Salud “Dr. Carlos G. Malbrán” (ANLIS-Malbrán, Argentina). Before each experiment, yeast strains were cultured on Sabouraud dextrose agar (SDA) at 37 °C for 24–48 h, as previously described [47].

### 2.3 Preparation of crude pepper seed extract

Crude extracts were prepared from *C. annuum* seeds using a modified plant protein-extraction protocol [48]. Briefly, approximately 30 g of dried seeds were ground into a fine powder using a mechanical blender. The resulting powder was suspended in 100 mL of 10 mM phosphate buffer (pH 7.4) and homogenized on ice to minimize oxidative browning and preserve extract integrity.

The homogenate was incubated at 4 °C for 3 h with agitation at 150 rpm and subsequently filtered through three layers of gauze to remove coarse particulate material. The filtrate was centrifuged at 7,000 × *g* for 30 min at 4 °C. The resulting supernatant, designated red bell pepper crude extract (RBPCE), was collected and stored at −20 °C until use.

### 2.4 Enrichment of thermostable low-molecular-weight components

Heat treatment was used as an efficient purification step to remove heat-labile components and enrich the extract in low-molecular-weight, thermostable nsLTPs, a class of proteins previously associated with antifungal activity [36,39,46]. The procedure was adapted from previously described thermal-purification protocols for *C. annuum* seed protease inhibitors [49].

Briefly, 1-mL aliquots of RBPCE were heated at 100 °C for 5 min in a water bath and subsequently centrifuged at 10,000 × *g* for 30 min at 4 °C to remove denatured and precipitated material. The resulting soluble supernatant was collected and designated red bell pepper heat-treated extract (RBPHT). Enrichment of low-molecular-weight, thermostable nsLTPs in RBPHT was subsequently assessed by Tricine–SDS–PAGE and LC-MS/MS analysis.

### 2.5 Protein concentration determination

The soluble protein concentration of RBPCE and RBPHT was determined using the Bradford assay, with BSA as the calibration standard [50]. A standard curve covering 0–1 mg/mL BSA was prepared, and 20 µL of each extract or standard was mixed with 200 µL of Bradford reagent in a 96-well microplate. After incubation for 10 min at room temperature, absorbance was measured at 595 nm using an Infinite M200 PRO microplate reader (Tecan, Männedorf, Switzerland). Measurements were performed in three technical replicates across four independent extract preparations.

### 2.6 Tricine–SDS–PAGE

Tricine–SDS–PAGE was used to compare the protein profiles of RBPCE and RBPHT and assess the enrichment of low-molecular-weight proteins following thermal treatment [51]. Samples were mixed with loading buffer containing 0.13 M Tris-HCl (pH 6.8), 2% (w/v) SDS, 5% (v/v) β-mercaptoethanol, 8% (v/v) glycerol, and 0.002% (w/v) bromophenol blue. Samples were heated at 100 °C for 5 min and subjected to electrophoresis using a Mini-PROTEAN III system (Bio-Rad, Hercules, CA, USA) at a constant current of 20 mA per gel. The three-part gels comprised a 4% stacking gel, a 10% spacer gel, and a 16% resolving gel. Following electrophoresis, proteins were stained with 0.2% (w/v) Coomassie Brilliant Blue G-250, and the gels were destained with distilled water.

### 2.7 In-gel digestion and LC–MS/MS protein identification

#### 2.7.1 Sample preparation

Gel bands were destained with 100 mM NH_4_HCO_3_ in 40% acetonitrile, reduced with dithiothreitol (2 µmol, 30 min, 56 °C), alkylated in the dark with iodoacetamide (11 µmols, 30 min, 25 °C), dehydrated with acetonitrile for trypsin digestion (400 ng, 37 °C for 8h, Promega cat # V5113). After digestion, peptide mix was acidified with formic acid and desalted with a MicroSpin C18 column (The Nest Group, Inc) prior to LC-MS/MS analysis.

#### 2.7.2 Chromatographic and MS analysis

Samples were analyzed using an Orbitrap Eclipse mass spectrometer (Thermo Fisher Scientific, San Jose, CA, USA) coupled to an EASY-nLC 1200 (Thermo Fisher Scientific (Proxeon), Odense, Denmark). Peptides were loaded directly onto the analytical column and were separated by reversed-phase chromatography using a 50-cm column with an inner diameter of 75 μm, packed with 2 μm C18 particles (Thermo Fisher Scientific, cat # ES903). Chromatographic gradients started at 95% buffer A and 5% buffer B with a flow rate of 300 nL/min and gradually increased to 25% buffer B and 75% A in 52 min and then to 40% buffer B and 60% A in 8 min. After each analysis, the column was washed for 10 min with 100% buffer B. Buffer A: 0.1% formic acid in water. Buffer B: 0.1% formic acid in 80% acetonitrile.

The mass spectrometer was operated in positive ionization mode with nanospray voltage set at 2.4 kV and source temperature at 305 °C. The acquisition was performed in data-dependent adcquisition (DDA) mode and full MS scans with 1 micro scans at resolution of 120,000 were used over a mass range of m/z 350-1400 with detection in the Orbitrap mass analyzer. Auto gain control (AGC) was set to ‘standard’ and injection time to ‘auto’. In each cycle of data-dependent acquisition analysis, following each survey scan, the most intense ions above a threshold ion count of 10,000 were selected for fragmentation. The number of selected precursor ions for fragmentation was determined by the “Top Speed” acquisition algorithm and a dynamic exclusion of 60 seconds. Fragment ion spectra were produced via high-energy collision dissociation (HCD) at normalized collision energy of 28% and they were acquired in the ion trap mass analyzer. AGC and injection time were set to ‘Standard’ and ‘Dynamic’, respectively and isolation window of 1.4 m/z was used.

Digested bovine serum albumin (New england biolabs cat # P8108S) was analyzed between each sample to avoid sample carryover and to assure stability of the instrument and Qcloud [52] has been used to control instrument longitudinal performance during the project.

#### 2.7.3 Data analysis

Acquired spectra were analyzed using the Proteome Discoverer software suite (v3.3, Thermo Fisher Scientific) and the Mascot search engine (v2.6, Matrix Science [53]). The data were searched against Uniprot *C. Annuum* database (as in October 2026), plus a list [54] of common contaminants and all the corresponding decoy entries. For peptide identification a precursor ion mass tolerance of 7 ppm was used for MS1 level, trypsin was chosen as enzyme and up to three missed cleavages were allowed. The fragment ion mass tolerance was set to 0.5 Da for MS2 spectra. Oxidation of methionine and N-terminal protein acetylation were used as variable modifications whereas carbamidomethylation on cysteines was set as a fixed modification. False discovery rate (FDR) in peptide identification was set to a maximum of 1%.

### 2.8 Trypsin inhibition assay

Trypsin inhibitory activity was determined using BApNA as the chromogenic substrate, following a modified version of the method described by Erlanger *et al*. [55]. Trypsin at a final concentration of 0.25 mg/mL was preincubated with increasing concentrations of RBPCE and RBPHT samples ranging from 0 to 2,000 µg/mL in 100 mM Tris-HCl buffer (pH 7.5). Extract concentrations were expressed as soluble protein. After incubation for 10 min at 37 °C, the reaction was initiated by adding BApNA to a final concentration of 1 mM [56]. All assays were performed in triplicate.

Trypsin inhibitory activity was expressed as the percentage reduction in proteolytic activity relative to the extract-free control. The half-maximal inhibitory concentration (IC₅₀), defined as the extract concentration required to reduce trypsin activity by 50%, was calculated from concentration–response curves by nonlinear regression fitting using GraphPad Prism v. 10.0.0 software (www.graphpad.com) [57].

### 2.9 Agar diffusion antifungal assay

The antifungal activity of RBPCE and RBPHT against *C. albicans*, *C. krusei*, and *C. auris* isolates belonging to clades I, III, and IV was evaluated using an agar diffusion assay based on the Kirby–Bauer method with minor modifications [58]. Yeast cultures were grown on SDA at 37 °C for 48 h. Cell suspensions were prepared in sterile physiological saline and adjusted to a 0.3 McFarland standard, corresponding to approximately 1 x 10^6^ cells/mL [59]. Aliquots of each suspension were spread uniformly across the surface of SDA plates.

To perform the assays, 200-µL of each extract was applied over the previously inoculated Sabouraud agar plates. Once the drop placed in the plate was dried, the plates were then incubated at 37 ° C for 24 h to allow fungal growth. After incubation diameters of the growth-inhibition zones were measured. In all cases, sterile extraction buffer was used as the negative control. Assays were performed in triplicate.

Commercial discs containing fluconazole (25 µg/disc), caspofungin (5 µg/disc), nystatin (50 µg/disc), or amphotericin B (10 µg/disc) were used as positive controls. The inoculation and incubation conditions were identical to those described for the extract diffusion assays. Following incubation at 37 °C for 24 h, inhibition-zone diameters were measured in millimetres.

### 2.10 Liquid-medium antifungal assays

#### 2.10.1. Determination of the minimum inhibitory and minimum fungicidal concentrations

The antifungal activity of RBPCE and RBPHT against a clade IV *C. auris* isolate was further evaluated by determining their minimum inhibitory concentration (MIC) and minimum fungicidal concentration (MFC). Yeast suspensions were prepared at a final density of 1 × 10^6^cells/mL and dispensed into 96-well microplates containing Sabouraud dextrose broth and increasing concentrations of each extract, in a final volume of 150 µL per well. Extract-free cultures and uninoculated medium containing the corresponding extract concentrations were included as growth controls and blanks, respectively. Plates were incubated at 37 °C, and optical density at 620 nm was recorded between 0 and 46 h using an Infinite M200 PRO microplate reader. All assays were performed in quadruplicate.

The minimum inhibitory concentration (MIC) was defined as the lowest extract concentration that completely inhibited visible fungal growth after 46 h of incubation. [60]. To determine the MFC, aliquots from wells showing at least 50% growth inhibition were plated onto Mueller–Hinton (MH) agar and incubated at 37 °C for 20 h. The MFC was defined as the lowest extract concentration that reduced the viable-cell count by at least 99.9%, corresponding to a reduction of ≥3 log₁₀ CFU relative to the initial inoculum [61]. Fluconazole was included as a reference antifungal drug. Yeast suspensions of the clade IV *C. auris* isolate were prepared at a final density of 1 × 10^6^ cells/mL and dispensed into 96-well microplates containing Sabouraud dextrose broth and fluconazole at a final concentration of 25 µg/mL. Untreated cultures and uninoculated medium were included as controls. Plates were incubated at 37 °C, and optical density at 620 nm was recorded between 0 and 46 h using an Infinite M200 PRO microplate reader.

### 2.11 Effects of RBPHT on biofilm development

The effects of RBPHT on planktonic-cell viability, initial adhesion, and biofilm development were evaluated against *C. albicans* and a clade IV *C. auris* isolate using 96-well microplates. Isolated colonies from 12–24-h cultures were inoculated into Sabouraud dextrose broth and incubated at 37 °C with agitation at 200 rpm until the exponential growth phase was reached. Cell suspensions were then adjusted to a final density of 1 × 10^6^ CFU/mL.

Two experimental conditions were evaluated (See experimental scheme in **Figure S2**). (I) To assess inhibition of initial yeast adhesion, wells were inoculated at time zero with the yeast suspension and a 3:1 (v/v) mixture of culture medium and RBPHT. After incubation for 1 h, the supernatant was removed and replaced with 150 µL of fresh, inoculum-free medium containing RBPHT at the same ratio (**Figure S2A**). (II) To assess inhibition of biofilm development, wells were initially inoculated in extract-free culture medium and incubated for 1 h to allow cell adhesion. The medium containing non-adherent cells was then removed and replaced with 150 µL of a 1:1 (v/v) mixture of culture medium and RBPHT (**Figure S2B**). Reference-antifungal controls were prepared following the same procedure as condition II, with RBPHT replaced by nystatin or fluconazole at final concentrations of 50 µg/mL and 25 µg/mL, respectively. Untreated wells containing the yeast suspension and culture medium served as biofilm-development controls, with the medium renewed after the initial 1-h adhesion period. Corresponding uninoculated blanks containing culture medium alone, medium plus RBPHT, or medium plus each reference antifungal were included to correct for background absorbance.

In all conditions, the supernatants removed after the initial 1-h incubation were retained for quantification of non-adherent viable cells. Samples were serially diluted in sterile physiological saline, plated onto Sabouraud dextrose agar, and incubated at 37°C for 48 hours. After a total incubation period of 24 h, supernatants containing planktonic cells were collected, and optical density was measured at 600 nm after subtraction of the corresponding blank. To quantify viable biofilm-associated cells, the wells were washed three times with sterile physiological saline to remove non-adherent cells. The attached biomass was then detached by thoroughly scraping the bottom of each well with a sterile pipette tip in 150 µL of physiological saline. The resulting suspensions were serially diluted and plated onto Sabouraud dextrose agar for CFU quantification. Each experimental condition was established in 12 parallel wells, which were combined into a representative pooled sample. Each pooled sample was serially diluted and plated for viable-cell quantification. The complete experiment was independently performed in triplicate.

### 2.12. Statistical analysis

Statistical analyses were performed using GraphPad Prism version 9.5.1 for Windows (GraphPad Software, San Diego, CA, USA; www.graphpad.com). Data are presented as the mean ± standard deviation (SD) or mean ± standard error of the mean (SEM), as indicated in the corresponding figure legends. Differences among multiple groups were analyzed using one-way or two-way analysis of variance (ANOVA), as appropriate, followed by Tukey’s multiple-comparisons post hoc test. Statistical significance was defined as *p* < 0.05.

## 3. Results and discussion

### 3.1 Thermal enrichment and identification of *C. annuum* seed peptides

Red bell pepper is widely consumed worldwide, both as a fresh vegetable and as a processed food product [62]. Beyond its nutritional value, *Capsicum* species have long been used in traditional medicine across different cultures to treat infections, inflammation, and skin disorders [63]. In recent years, biochemical studies have identified several small, cysteine-rich peptides in *C. annuum* tissues, including seed-derived members of the nsLTP family with antifungal activity against pathogenic yeasts [36,39,46]. More recently, an nsLTP2 isolated from *C. annuum* leaves was also shown to inhibit the growth and reduce the viability of several fungal pathogens, including *C. albicans* [38]. We previously isolated a multifunctional thermostable AMP from yellow *C. annuum* seeds using heat treatment as an initial purification step [49].

Building on these findings, we applied a similar thermal-fractionation strategy to red *C. annuum* seeds to enrich the thermostable peptides present in this material and evaluate its antifungal activity against *C. auris*. Extraction under mild conditions yielded a crude red bell pepper seed extract (RBPCE) with a soluble protein concentration of approximately 6.0 mg/mL (**Figure 1A**). RBPCE was subsequently heated at 100 °C for 5 min and centrifuged to remove denatured and precipitated material. The resulting soluble heat-treated fraction, designated RBPHT, contained approximately 1.5 mg/mL soluble protein.

**Figure 1.**
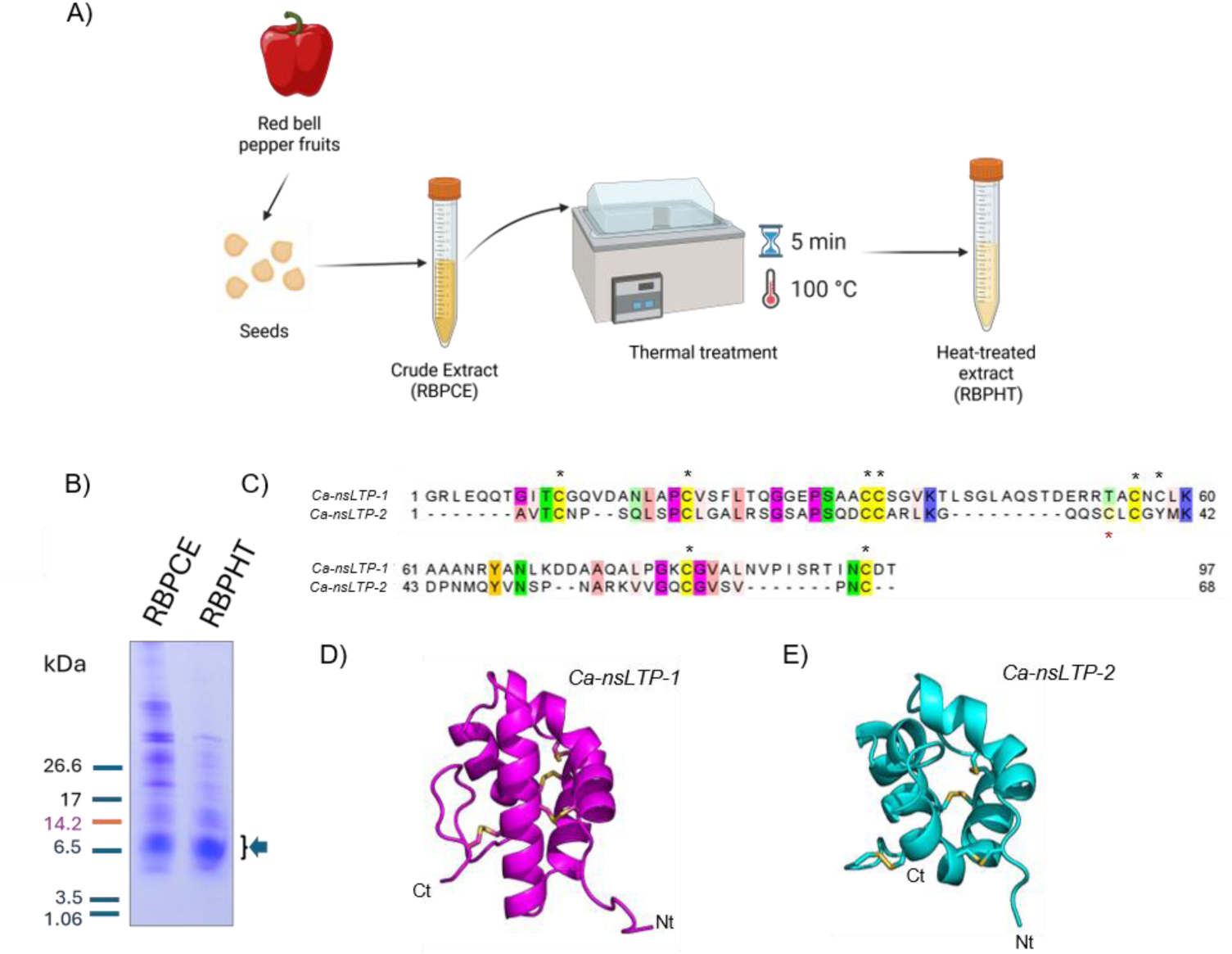
Isolation and characterization of thermostable pepper seed nsLTPs-enriched fraction. (A) Schematic representation of the extraction and thermal-fractionation procedure used to obtain the crude red bell pepper seed extract (RBPCE) and the heat-treated, nsLTP-enriched fraction (RBPHT). (B) Tricine–SDS–PAGE profiles of RBPCE and RBPHT, showing the enrichment of low-molecular-weight proteins following heat treatment. The arrow indicates the major RBPHT band, with an apparent molecular mass of approximately 8 kDa. Molecular-mass markers are indicated in kDa. (C) Sequence alignment of the mature Ca-nsLTP1 and Ca-nsLTP2 proteins identified by LC–MS/MS analysis of the major RBPHT peptide components. Cysteine residues involved in disulfide-bond formation are indicated by asterisks. (D, E) AlphaFold-predicted three-dimensional structures of Ca-nsLTP1 (D) and Ca-nsLTP2 (E), corresponding to type I and type II nsLTPs, respectively. Their disulfide-bond pairings are C1–C6, C2–C3, C4–C7 and C5–C8 for Ca-nsLTP1, and C1–C5, C2–C3, C4–C7 and C6–C8 for Ca-nsLTP2. The N and C termini are labelled Nt and Ct, respectively. Panel A was created with BioRender.com. Structural models were generated using AlphaFold [64].

Tricine–SDS–PAGE showed that thermal fractionation markedly simplified the protein profile and enriched low-molecular-weight components migrating below 15 kDa (**Figure 1B**). A major band with an apparent molecular mass of approximately 8 kDa accounted for approximately 75% of the combined densitometric intensity of these bands. LC–MS/MS analysis identified Ca-nsLTP1 (A0A2G3AMC5) and Ca-nsLTP2 (A0A2G3AAS4) as the predominant protein constituents detected in this fraction, with sequence coverages of 51% and 47% of the predicted mature proteins, respectively (**Table S1**). Their calculated mature molecular masses, approximately 9.5 and 7.1 kDa, are consistent with their co-migration within the major RBPHT band. Detailed sequence and structural analyses indicated that Ca-nsLTP1 and Ca-nsLTP2 belong to the type I and type II nsLTP subfamilies, respectively, each displaying the characteristic cysteine connectivity and compact α-helical architecture [65,66] (**Figure 1C–E**). The identification of these proteins in *C. annuum* seeds is consistent with previous reports of related nsLTPs in different tissues of this plant. Ca-nsLTP1 closely matches the experimentally determined N-terminal sequence of Ca-LTP1, a 9-kDa nsLTP previously isolated from *C. annuum* seeds by Cruz *et al.* [39] and subsequently characterized in greater detail by Diz et al. [36], sharing 21 of 23 unambiguous aligned residues. Similarly, the 33-residue C-terminal sequence reported for the leaf-derived type II protein CaCLTP2 is identical to the corresponding region of Ca-nsLTP2, although the latter contains an additional four-residue insertion (DPNM) [38].

Protease-inhibitory activity has also been reported for different plant nsLTPs [67,68]. As shown in **Figure S1**, both RBPCE and RBPHT strongly inhibited trypsin, reaching approximately 96% and 98% inhibition, respectively, at the highest concentrations tested (1,520 µg/mL for RBPCE and 640 µg/mL for RBPHT). RBPHT exhibited a substantially lower IC₅₀ than RBPCE (290 ± 4 versus 800 ± 6 µg/mL), corresponding to an approximately 2.8-fold increase in inhibitory potency on a soluble-protein basis. Thus, thermal fractionation retained and enriched trypsin-inhibitory activity within the soluble nsLTP-enriched fraction. This finding is also consistent with our previous isolation of the thermostable trypsin inhibitor (YBPTI) from yellow *C. annuum* seeds using an equivalent heat-treatment step [49]. In a similar manner, a type I nsLTP isolated from *Ginkgo biloba* seeds inhibited papain and pepsin and produced inhibition of trypsin under the tested conditions [68]. In a more recent study, an nsLTP purified from *Eleusine coracana* seeds was reported to possess dual α-amylase-and trypsin-inhibitory activities [67]. Interestingly, sequence and structural comparisons showed that Ca-nsLTP1 shares a conserved compact α-helical fold with these proteins, with RMSD values of 0.767 Å relative to the *E. coracana* nsLTP and 0.323 Å relative to *G. biloba* nsLTP1 (**Figure S1**).

Overall, thermal fractionation yielded a thermostable, low-molecular-weight fraction enriched in type I and type II nsLTPs. This enrichment coincided with increased trypsin-inhibitory potency, although the specific contribution of Ca-nsLTP1 to this activity requires further investigation. Given the previously reported antifungal properties of related nsLTPs, we next evaluated the activity of RBPHT against clinically relevant *Candida* species, including the emerging pathogen *C. auris*.

### 3.2. The nsLTP-enriched fraction exhibits antifungal activity against multiple *C. auris* clades

The antifungal activity of RBPHT was initially evaluated using agar diffusion assays against a panel of clinically relevant fungal strains, comprising *C. albicans*, *C. krusei*, and *C. auris* isolates belonging to clades I, III, and IV (**Figure 2** and **Table 1**).

**Figure 2.**
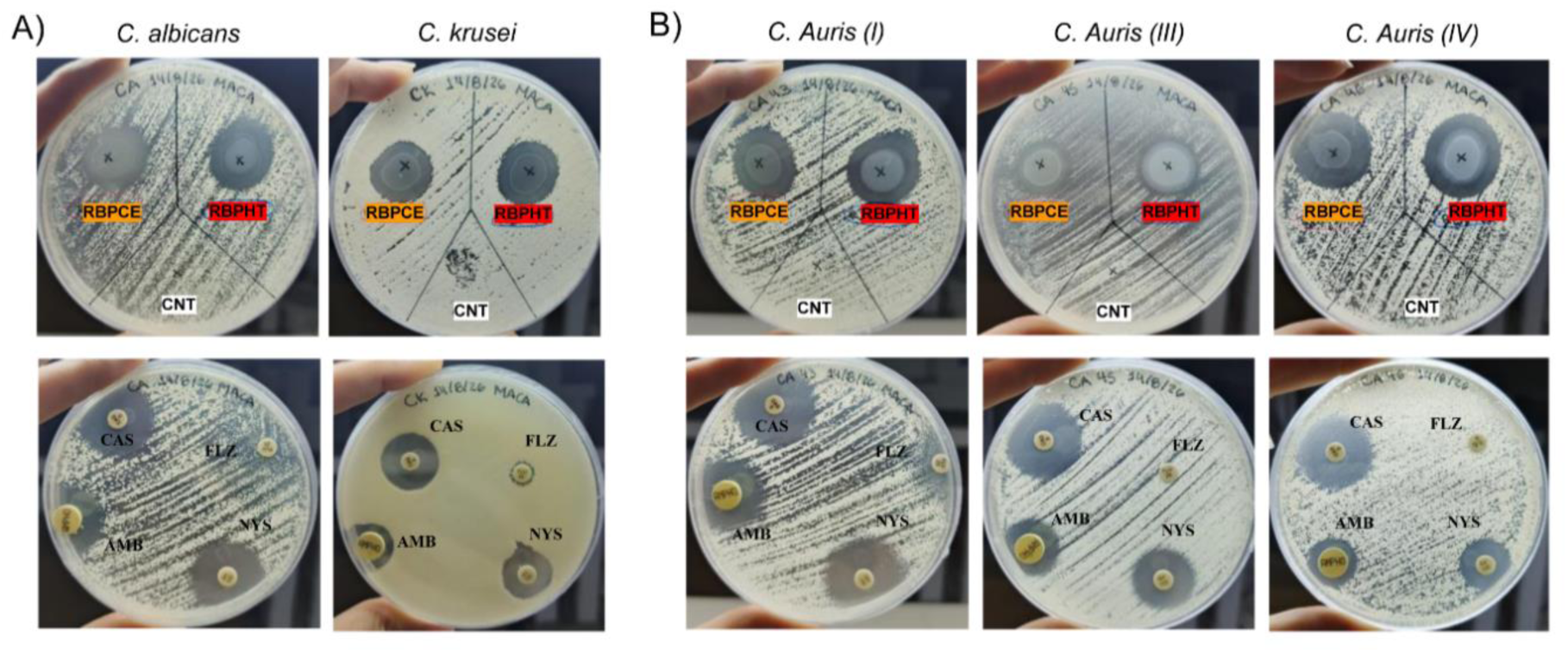
Antifungal activity of the nsLTP-enriched RBPHT fraction against pathogenic fungal strains. (A) Representative agar diffusion assays showing the activity of RBPHT against *C. albicans* and *C. krusei*. The untreated crude extract (RBPCE) was included as the starting-extract reference. (B) Activity of RBPHT and RBPCE against *C. auris* isolates belonging to clades I, III, and IV. Both preparations produced measurable inhibition zones against all strains tested, whereas the phosphate extraction buffer used as a negative control (CNT) produced no detectable inhibition. The lower panels show the inhibition zones produced by the reference antifungal agents fluconazole (FLZ; 25 μg), caspofungin (CAS; 5 μg), nystatin (NYS; 50 μg), and amphotericin B (AMB; 10 μg). No fluconazole inhibition zone was detected for the clade III or IV *C. auris* isolates.

**Table 1.**
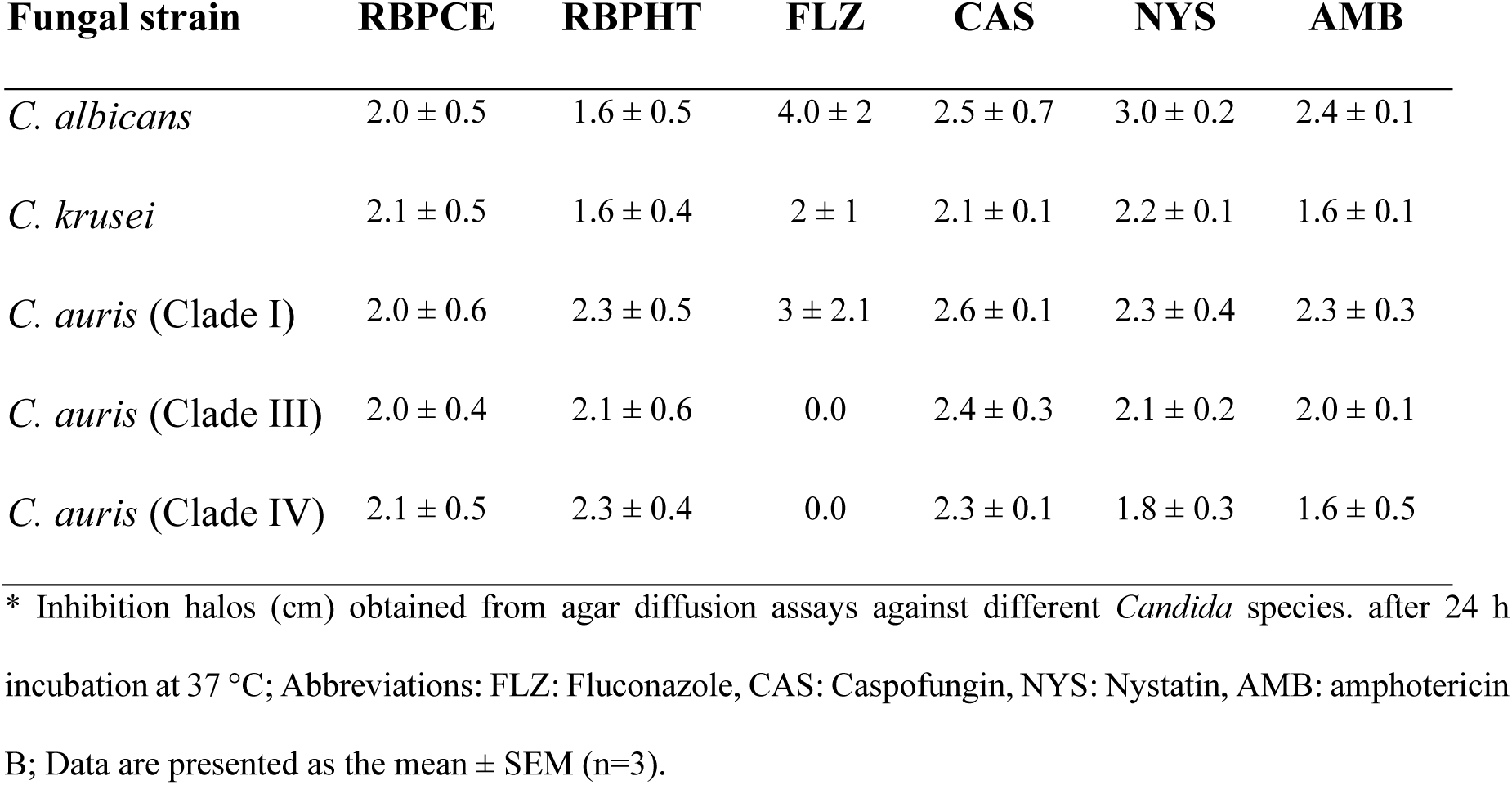
Inhibition-zone diameters produced by RBPCE, RBPHT, and reference antifungal agents against *Candida* and *Candidozyma* strains.

RBPHT inhibited all five strains tested, producing inhibition zones of 1.6 cm against *C. albicans* and *C. krusei* and values ranging from 2.1 to 2.3 cm against the three *C. auris* isolates (**Table 1**). RBPCE also inhibited all strains and produced zones ranging from 2.0 to 2.1 cm. Thus, although thermal treatment reduced the soluble protein concentration by approximately 75%, RBPHT retained antifungal activity comparable to that of the untreated starting extract and showed no apparent reduction in activity against *C. auris*. No inhibition was observed with the phosphate extraction buffer.

Comparison with the reference antifungal agents revealed differences among the strains tested (**Figure 2** and **Table 1**). Caspofungin, nystatin, and amphotericin B produced measurable inhibition zones against all strains. Fluconazole produced inhibition zones against *C. albicans*, *C. krusei*, and the clade I *C. auris* isolate, whereas no detectable zone was observed for the clade III or IV isolates, consistent with their reduced fluconazole susceptibility [7]. More importantly, RBPHT inhibited all three *C. auris* isolates irrespective of their response to fluconazole, indicating that its activity was retained across genetically distinct clades and was not associated with azole susceptibility.

The activity of RBPHT against fluconazole-resistant *C. auris* isolates, together with its enrichment in Ca-nsLTP1 and Ca-nsLTP2, supports further investigation of these thermostable nsLTPs as candidate antifungal peptides against this pathogen.

### 3.3. Concentration-Dependent Effects of nsLTP-enriched RBPHT fraction on *C. auris* Growth Kinetics

#### 3.3.1. Effects of RBPHT on C. auris growth kinetics

To further characterize the antifungal activity of the pepper seed extracts quantitatively, we examined their concentration-dependent effects on the growth kinetics of a representative clade IV *C. auris* isolate. This isolate belongs to the South American lineage and displayed no detectable inhibition zone in response to fluconazole in the agar diffusion assay described above (**Figure 2)**.

In untreated cultures, *C. auris* exhibited a typical growth profile, characterized by a lag phase followed by a marked increase in optical density (**Figure 3A**). RBPCE inhibited fungal growth in a concentration-dependent manner. Concentrations between 190 and 390 μg/mL delayed growth and reduced the final culture density, whereas complete suppression of detectable growth was observed at concentrations of 780 μg/mL or higher. The nsLTP-enriched RBPHT fraction exhibited greater inhibitory activity on a soluble-protein concentration basis: concentrations between 90 and 180 μg/mL markedly reduced fungal growth, and no measurable increase in optical density was observed at concentrations of 180 μg/mL or higher. Fluconazole at 25 μg/mL was included as a reference antifungal control.

**Figure 3.**
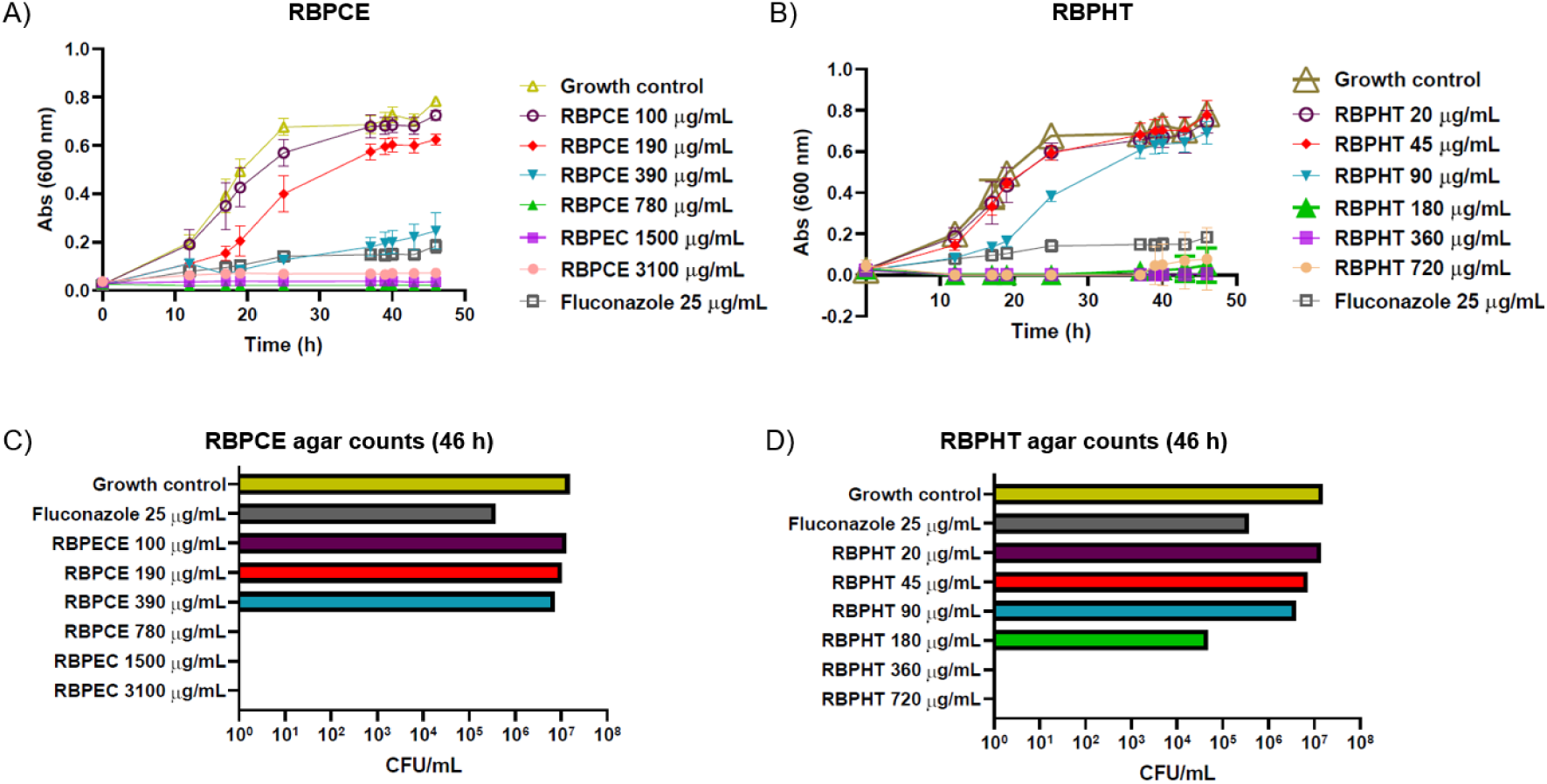
Concentration-dependent effects of *C. annuum* nsLTP-enriched RBPHT fraction on *C. auris* growth and viability. (A, B) Growth kinetics of a clade IV *C. auris* isolate cultured in liquid medium in the absence or presence of increasing concentrations of (A) RBPCE or (B) RBPHT, added at the beginning of the experiment (t = 0 h). Untreated cultures and cultures containing fluconazole (25 μg/mL) were included as controls. In all cases, data are presented as the mean ± SD (n=4). (C,D) Viable-cell counts were determined after 46 h of incubation with (C) RBPCE or (D) RBPHT.

Viable-cell counts obtained after 46 h were consistent with the growth-curve results (**Figure 3C** and **3D**). Both extracts reduced the number of recoverable viable cells in a concentration-dependent manner, with nsLTP-enriched RBPHT fraction producing a stronger effect at lower concentrations than RBPCE. These results demonstrate that the antifungal activity present in the crude extract was retained and substantially enriched on a soluble-protein concentration basis, following heat treatment.

#### 3.3.2. Determination of MIC and MFC values

Using the absence of detectable fungal growth as the endpoint, RBPHT exhibited an MIC of 180 µg/mL against the clade IV *C. auris* isolate, compared with 780 µg/mL for the untreated RBPCE reference (**Figures 3A and 3B**). Thus, RBPHT showed an approximately 4.3-fold lower MIC than the starting crude extract. Subculturing cells from wells without detectable growth onto extract-free agar yielded MFC values of 360 µg/mL for RBPHT and 780 µg/mL for RBPCE (**Figures 3C and 3D**). The corresponding MFC/MIC ratios were 2 and 1, respectively, indicating fungicidal activity under the conditions tested. Collectively, these results demonstrate that thermal fractionation increased both the growth-inhibitory and fungicidal potency of the soluble fraction.

For context, the tentative fluconazole-resistance breakpoint proposed for *C. auris* is ≥32 µg/mL, and MIC values ≥256 µg/mL are frequently observed among resistant isolates. Nevertheless, the concentrations determined for RBPHT and fluconazole should not be compared directly. Fluconazole is a defined low-molecular-weight compound, whereas RBPHT consists predominantly of macromolecular proteinaceous components and is expressed in terms of total protein mass.

The increased antifungal potency of RBPHT is consistent with its enrichment in Ca-nsLTP1 and Ca-nsLTP2. Several studies have demonstrated antifungal activity for related *Capsicum* nsLTPs. A peptide fraction containing an approximately 9-kDa seed-derived nsLTP inhibited the growth of *C. albicans* and *C. tropicalis* [39]. Subsequent characterization of the purified protein, designated Ca-LTP1, confirmed its activity against *C. tropicalis* and demonstrated plasma-membrane permeabilization, supporting membrane disruption as a possible mechanism of action [36]. In a recent study, LTP-enriched fractions obtained from *C. chinense* seeds inhibited *C. albicans* growth and produced predominantly fungistatic effects [46]. The type II protein CaCLTP2, isolated from *C. annuum* leaves, was also shown to inhibit several clinically relevant yeast species and induce membrane permeabilization, oxidative stress, metacaspase activation, and loss of cell viability [38]. Taken together, these findings suggest that Ca-nsLTP1 and Ca-nsLTP2 as plausible contributors to the antifungal activity of RBPHT, although the individual effects of these AMPs against *C. auris* remain to be established using the isolated proteins.

We previously purified the thermostable trypsin inhibitor YBPTI from yellow *C. annuum* seeds and demonstrated fungicidal activity against *C. albicans*, with an MIC of 5.7 µg/mL, together with disruption of plasma-membrane integrity [49]. This observation is relevant because RBPHT retained strong trypsin-inhibitory activity following thermal peptide enrichment. Accordingly, the thermostable inhibitors retained in the nsLTP-enriched RBPHT fraction may reduce *C. auris* growth and viability by interfering with protease-dependent processes and/or compromising fungal membrane function [30,69].

Taken together, these results show that the nsLTP-enriched RBPHT fraction induces concentration-dependent growth inhibition and loss of viability in a clade IV *C. auris* isolate. Several plant-derived AMPs have been shown to target *Candida* biofilms at different stages. The plant defensin HsAFP1 inhibits *C. albicans* biofilm formation and acts synergistically with caspofungin against established biofilms [70], whereas the *Arabidopsis thaliana*-derived decapeptide OSIP108 interferes with biofilm development without affecting planktonic growth or viability [71]. More recently, the barley defensin D-lp1 was shown to inhibit *C. auris* adhesion and biofilm formation and to disrupt mature biofilms [72]. In this context, we next evaluated the potential of the nsLTP-enriched RBPHT fraction to target *C. auris* adhesion and biofilm development.

### 3.4 The nsLTP-enriched RBPHT fraction inhibits *C. auris* biofilm development

Beyond planktonic growth, surface adhesion and biofilm formation are important virulence attributes of pathogenic fungal species, contributing to their persistence on host tissues, medical devices, and environmental surfaces [11,14]. Biofilm development generally proceeds through sequential stages comprising initial adhesion, proliferation and microcolony formation, extracellular matrix production, maturation, and the eventual dispersal of cells capable of colonizing new sites (**Figure 4A**). Although biofilm architecture varies among *Candida* and *Candidozyma*, cells embedded within mature biofilms generally exhibit reduced susceptibility to antifungal treatment, emphasizing the need for compounds capable of interfering with biofilm establishment and development [73,74].

**Figure 4.**
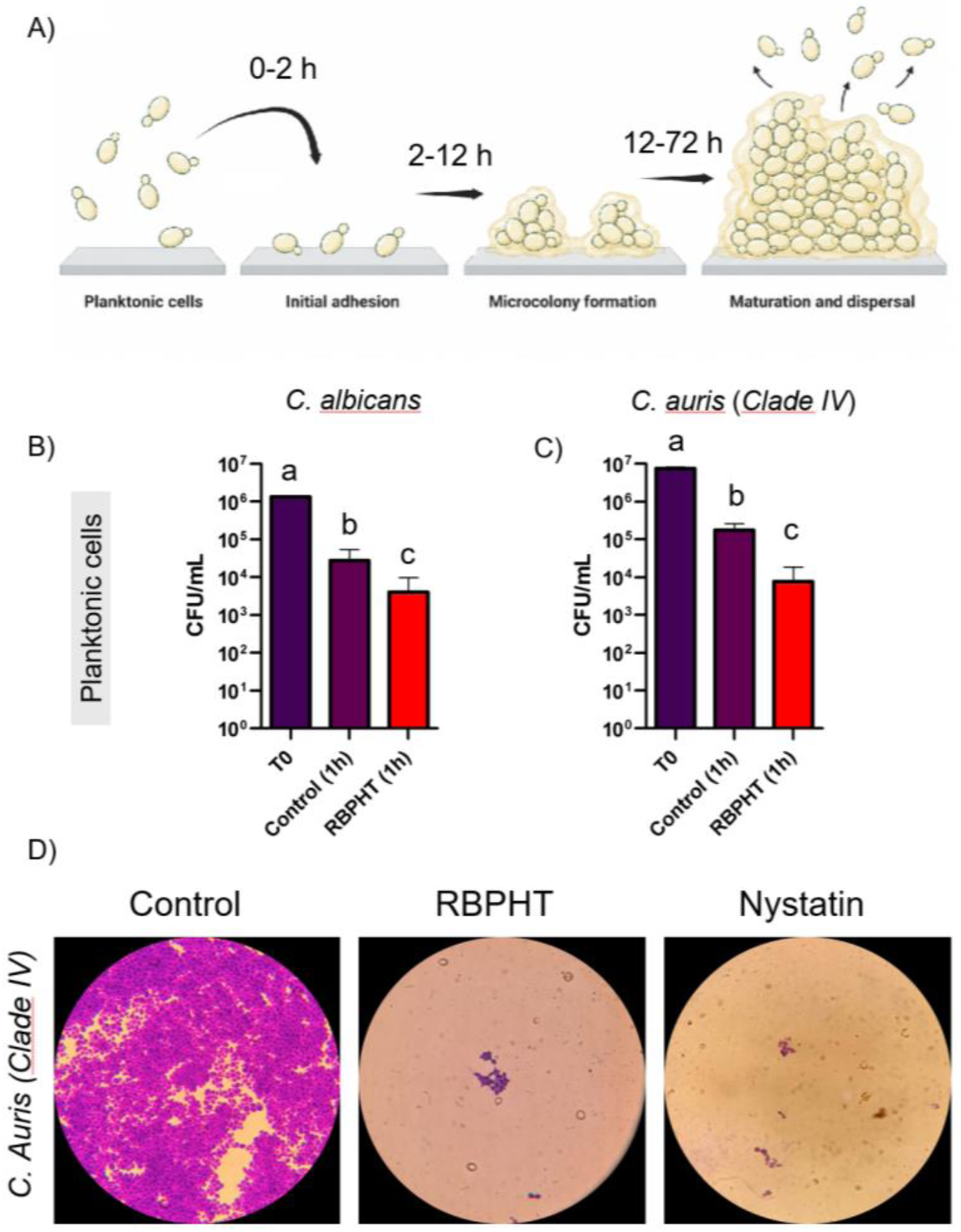
Effects of nsLTP-enriched RBPHT on planktonic-cell viability and biofilm development. (A) Schematic overview of *C. auris* biofilm development, comprising initial adhesion, microcolony formation, maturation, and cell dispersal; approximate developmental times are indicated. Created in https://BioRender.com. (B,C) Viable planktonic cells recovered at the beginning of the experiment (T0) and after the initial 1-h adhesion period in untreated and RBPHT-treated cultures of (B) *C. albicans* and (C) the clade IV *C. auris* isolate. Results are expressed as CFU/mL and presented as mean ± SD (n=3). Different lowercase letters indicate statistically significant differences between treatments (p < 0.05, one-way ANOVA followed by Tukey’s post hoc test). (D) Representative light microscopy images of 24 h *C. auris* (Clade IV) biofilms fixed on glass slides and stained with crystal violet. Control: untreated biofilm showing dense cellular aggregation; RBPHT: biofilm formed in the presence of RBPHT treatment; Nystatin: biofilm formed in the presence of nystatin. All images were captured using a 100x oil-immersion objective.

We therefore evaluated nsLTP-enriched RBPHT fraction against the clade IV *C. auris* isolate and included *C. albicans* as a well-characterized reference model. As illustrated in **Figure S2**, two complementary experimental conditions were examined: exposure to RBPHT from the time of inoculation to assess its effect on initial adhesion (**Figure S2A**), and addition of RBPHT after a 1-h adhesion period to assess its effect on subsequent biofilm development (**Figure S2B**). Planktonic and biofilm-associated cells were subsequently quantified to distinguish the effects of RBPHT on non-adherent cells, surface attachment, and the viability of the developing biofilm.

#### 3.4.1 nsLTP-enriched RBPHT suppresses early biofilm establishment

The effect of RBPHT during the initial adhesion phase was evaluated by exposing *C. auris* (clade IV) and *C. albicans* isolates to the extract from the time of inoculation (**Figure S2A**). In both pathogenic strains, the number of viable non-adherent cells recovered from the supernatant (SN1 in **Figure S2A**) decreased significantly after 1 h relative to the initial inoculum (T0), consistent with cell attachment to the well surface (**Figure 4B and 4C**) [75–77]. Notably, RBPHT treatment caused an additional significant reduction in viable non-adherent cells in both strains, compared with the corresponding untreated cultures after 1 h (**Figure 4B** and **4C**). This finding is consistent with the inhibitory effect of RBPHT on planktonic growth described above.

At 24 h post-inoculation, the supernatants (SN2 and SN3 in **Figure S2A**) were also collected for spectrophotometric analysis and plate counting. Additionally, the bottom of all wells was scraped to detach the already formed biofilm. The scraped biomass was resuspended in saline solution for viable cell quantification (SN3 in **Figure S2A**). Both quantifications revealed complete inhibition of biofilm formation or planktonic cells in the RBPHT-treated wells. By contrast, untreated biofilm-development controls contained approximately 1 x 10^7^ CFU/mL for *C. albicans* and 1 x 10^8^ CFU/mL for *C. auris*. The corresponding untreated planktonic populations reached 3.7 x 10^7^ and 1.4 x 10^7^ CFU/mL, respectively. Representative microscopy images similarly showed markedly less surface-associated *C. auris* biomass following treatment with RBPHT or nystatin than under untreated conditions (**Figure 4D**).

#### 3.4.2 RBPHT inhibits biofilm development after initial adhesion

Having demonstrated that exposure from the time of inoculation prevents biofilm establishment, we next examined whether nsLTP-enriched RBPHT could inhibit biofilm development after initial cell attachment. This question is clinically relevant because attached cells progressively form a protective extracellular matrix and acquire biofilm-associated phenotypes that reduce antifungal penetration and susceptibility, making established biofilms substantially more difficult to eradicate than planktonic populations [78,79].

Cells were allowed to adhere for 1 h in the absence of treatment, after which non-adherent cells were removed and the attached populations were incubated with RBPHT for 24 h (**Figure S2B**). Parallel cultures treated with FLZ or NYS were included as reference-antifungal controls, whereas untreated cultures served as biofilm-development controls.

RBPHT markedly reduced the number of viable biofilm-associated cells, producing an approximately 4-log₁₀ reduction in *C. albicans* and a greater than 2-log₁₀ reduction in *C. auris* relative to their respective untreated controls (**Figure 5A** and **5B**). Under the conditions tested, these reductions were greater than those produced by NYS, whereas FLZ had little or no detectable effect on the viable biofilm burden. These findings demonstrate that nsLTP-enriched RBPHT acts on previously attached cells and strongly inhibits their subsequent proliferation and development into viable biofilms.

**Figure 5.**
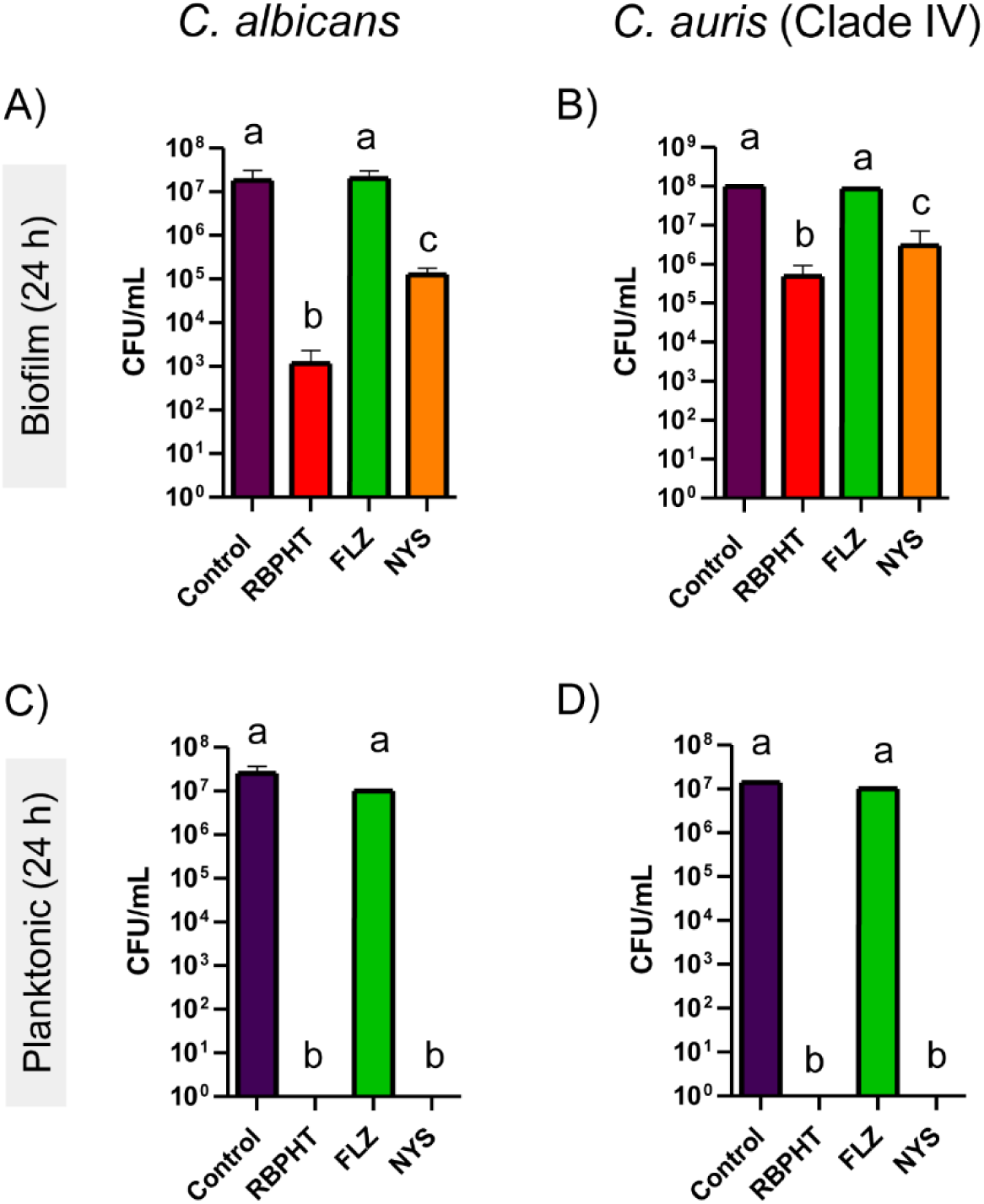
Effects of RBPHT on biofilm development and planktonic-cell viability after initial adhesion. Cells were allowed to adhere for 1 h before exposure to RBPHT or the reference antifungals. (A,B) Viable biofilm-associated cells recovered after 24 h from (A) *C. albicans* and (B) the clade IV *C. auris* isolate treated with RBPHT, fluconazole (FLZ), or nystatin (NYS). (C,D) Viable non-adherent cells recovered from the corresponding 24-h supernatants of (C) *C. albicans* and (D) *C. auris* cultures. Untreated cultures served as biofilm-development controls. Results are expressed as CFU/mL and presented as the mean ± SD (n=3). Within each panel, groups labelled with different lowercase letters differ significantly (*p* < 0.05; one-way ANOVA followed by Tukey’s multiple-comparisons test), whereas groups sharing the same letter are not significantly different.

As described above, the number of non-adherent cells initially decreased during the 1-h adhesion period. During subsequent biofilm development, however, viable cells in the supernatant may originate from both planktonic proliferation and the detachment or dispersal of biofilm-associated cells. In *C. albicans*, dispersal occurs throughout biofilm development and releases predominantly yeast-form cells, many of which originate through lateral budding from biofilm-associated hyphae [17,80]. By contrast, *C. auris* lacks the extensive hyphal network characteristic of *C. albicans* and exhibits both aggregative and non-aggregative phenotypes; accordingly, its non-adherent population may comprise individual yeast cells and small cellular aggregates [81–83].

No viable cells were recovered from the 24-h supernatants of RBPHT-treated cultures of either species, indicating that viable non-adherent populations were reduced below the detection limit of the assay (**Figure 5C** and **5D**). A similar effect was observed with NYS, whereas FLZ-treated cultures retained viable-cell counts comparable to those of the untreated controls. Together, these results demonstrate that RBPHT inhibits biofilm development from previously attached cells while suppressing the accumulation of viable cells in the surrounding liquid phase, potentially limiting biofilm expansion and colonization of new surfaces.

## 4. Conclusions

In this study, we demonstrate that a thermostable fraction derived from *C. annuum* seeds exhibits antifungal activity against multiple clinically relevant yeast species, including genetically distinct clades of the emerging multidrug-resistant pathogen *C. auris*. To our knowledge, this is the first report of antifungal activity against *C. auris* by a *Capsicum* seed-derived AMPs. Importantly, RBPHT inhibited isolates from three genetically distinct *C. auris* clades, including clade III and IV isolates that showed no detectable response to fluconazole under the conditions tested.

Thermal fractionation markedly simplified the protein profile while retaining and increasing the antifungal potency of RBPHT. LC–MS/MS analysis of its major electrophoretic band identified Ca-nsLTP1 and Ca-nsLTP2 as the predominant protein constituents. These type I and type II nsLTPs possess the compact, cysteine-rich architecture characteristic of thermostable plant defence proteins [41,43,65,66]. Thermal fractionation also enriched trypsin-inhibitory activity, although this activity cannot yet be attributed exclusively to either nsLTP. Similarly, the individual contributions of Ca-nsLTP1 and Ca-nsLTP2 to the antifungal activity of RBPHT remain to be precisely established.

Preventing biofilm establishment is particularly relevant to infections associated with biomaterials and catheters [1,14]. Owing to their high cellular density and protective extracellular matrix, mature biofilms may require antifungal concentrations substantially higher than those effective against planktonic cells [84,85]. Beyond inhibiting planktonic growth, RBPHT suppressed early biofilm establishment in both *C. albicans* and *C. auris*. The fraction prevented the recovery of viable biofilm-associated cells when present from inoculation and markedly reduced viable populations when added after initial cell adhesion, demonstrating activity against previously attached cells. Related *Capsicum* nsLTPs have been reported to permeabilize fungal plasma membranes and induce oxidative stress and loss of viability [37]. Such effects could reduce the viable founder population and prevent its progression into a mature, matrix-embedded community. Nevertheless, the present results do not establish a precise biofilm-specific mechanism, and the observed effects may arise, at least partly, from the fungicidal activity of RBPHT against both planktonic and attached cells.

Overall, these findings identify thermostable seed nsLTPs as emerging plant-derived antimicrobial peptides with antibiofilm properties and support their further exploration as candidates for targeting *C. auris* colonization and biofilm development. Their compact, disulfide-stabilized structures, together with their predominance in a fraction active against attached fungal cells, may be particularly relevant to strategies aimed at limiting fungal persistence on clinically relevant surfaces.

## ACKNOWLEDGEMENTS

J.G.-P. is associate professor at the Dpt. of Biochemistry and Molecular Biology from the Universitat Autònoma de Barcelona. W.D.O. is a member of the Researcher Career Program of the CONICET. S.M.G.V is a doctoral fellow from National Agency of Scientific and Technological Promotion ANPCyT (Argentina). The authors acknowledge the support from Universidad Nacional de La Plata (Argentina) under grants UNLP I+D 2020 project Code X-920 and UNLP I+D 2025 project Code X-1070; the ANPCyT under Project PICT-2021-00955; and PIP-CONICET under Project 815-2022-2024. The authors are grateful to Dr. Constanza Taverna from the National Institute of Infectious Diseases (INEI) - ANLIS ‘Dr. Carlos G. Malbrán’ (Buenos Aires, Argentina) for the generous donation of the *C. auris* clinical isolates used in this study. JGP was supported by the Spanish Ministry of Science and Innovation grant PID2025-174702OA-I00.

## AUTHOR CONTRIBUTIONS

S.M.V.G., C.L., S.C., J.G.-P. and W.D.O. performed experiments, analyzed the data, and wrote the initial draft of the manuscript. S.M.V.G, J.M.-R., J.G.-P. and W.D.O. assisted with the design of experiments and with the writing and revision of the manuscript. All authors have read and agreed to the published version of the manuscript.

## DECLARATION OF GENERATIVE AI AND AI-ASSISTED TECHNOLOGIES

The authors employed ChatGPT solely to enhance the readability and language quality of this manuscript. All content was subsequently reviewed and revised by the authors, who retain full accountability for the published work.

