## Supporting Information for "Antifungal and Antibiofilm Activity of a Thermostable Lipid Transfer Protein-Enriched Fraction from *Capsicum annuum* L. Seeds against *Candidozyma auris*"

### SUPPLEMENTARY FIGURES

#### Figure S1

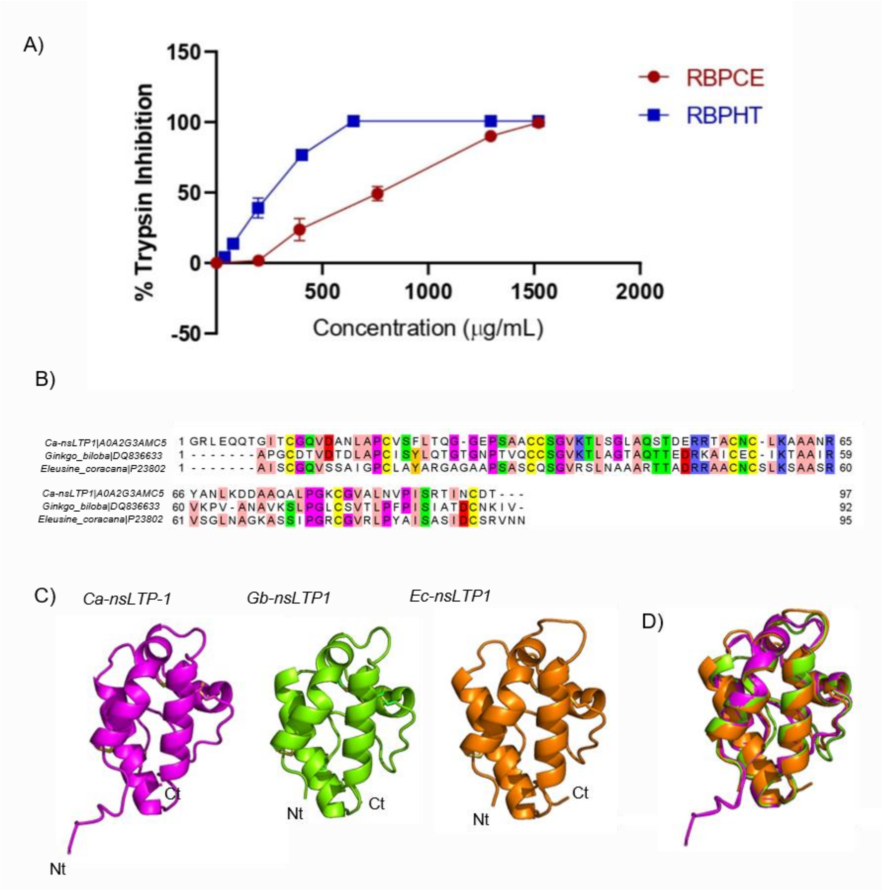


**Figure S1. Trypsin-inhibitory activity of pepper seed fractions and structural comparison of Ca-nsLTP1 with related plant proteins.** (A) Concentration-dependent inhibition of trypsin by the crude red bell pepper seed extract (RBPCE) and the heat-treated, nsLTP-enriched fraction (RBPHT). Values are presented as the mean ± SD (n=3). (B) Multiple-sequence alignment of mature Ca-nsLTP1 (A0A2G3AMC5), *Ginkgo biloba* nsLTP1 (DQ836633), and the nsLTP-like bifunctional α-amylase/trypsin inhibitor from *Eleusine coracana* (P23802). Conserved residues are highlighted. (C) Structural models of Ca-nsLTP1 (magenta), *G. biloba* nsLTP1 (green), and the *E. coracana* bifunctional inhibitor (orange), respectively. The N and C termini are labelled Nt and Ct. (D) Structural superposition of the three proteins. Relative to Ca-nsLTP1, the *G. biloba* and *E. coracana* proteins displayed RMSD values of 0.323 and 0.767 Å, respectively. Structural models were generated using AlphaFold [1].

**Figure S2**

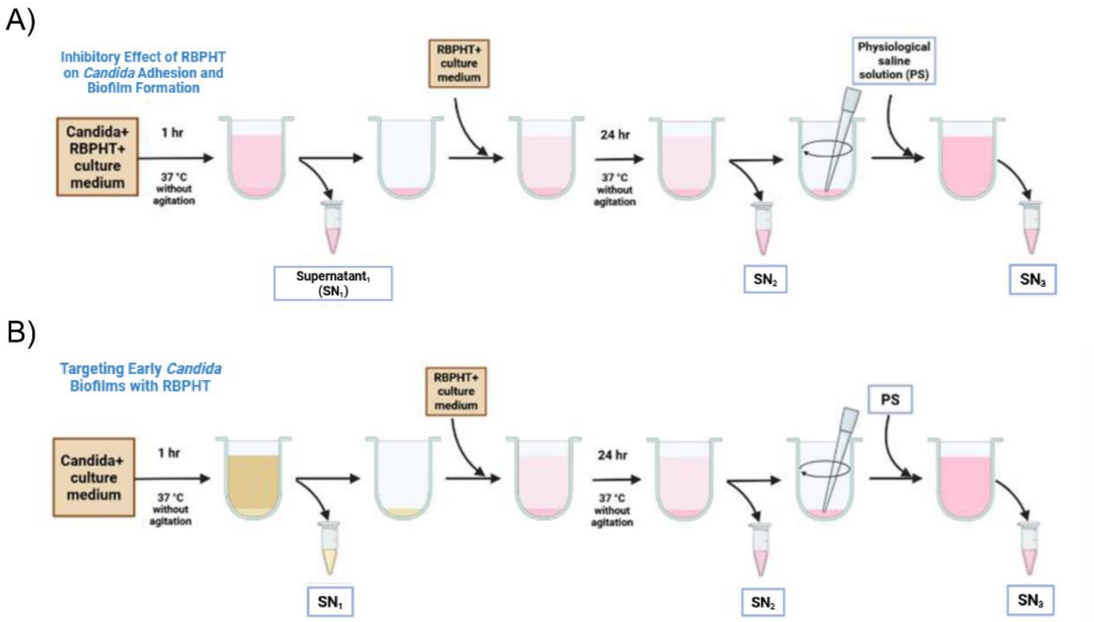


**Figure S2. Experimental design for evaluating the effects of RBPHT on yeast adhesion and biofilm development.** (A) To evaluate the effect of RBPHT on initial adhesion, *C. albicans* and the clade IV *C. auris* isolate were exposed to RBPHT from the time of inoculation. After 1 h, the supernatant containing non-adherent cells (SN1) was collected and replaced with fresh medium containing RBPHT, followed by incubation for 24 h. (B) To evaluate inhibition of biofilm development after initial cell attachment, cells were allowed to adhere for 1 h in the absence of RBPHT. The supernatant containing non-adherent cells (SN1) was then collected and replaced with fresh medium containing RBPHT, followed by incubation for 24 h. At the end of both assays, the 24-h culture supernatant containing non-adherent cells (SN2) was collected. The wells were subsequently washed, and biofilm-associated cells (SN3) were recovered by scraping into physiological saline (PS). Untreated biofilm-development controls and, where indicated, reference-antifungal controls containing nystatin or fluconazole were evaluated in parallel.

### SUPPLEMENTARY TABLES

#### Table S1. LC–MS/MS identification of major RBPHT proteins following in-gel tryptic digestion.

<table>
<tr><th>Protein</th><th>Accession</th><th>Protein name</th><th>MW (kDa)</th><th>MW w/o SP (kDa)</th><th>pI (mature)</th><th>Coverage (%) (mature)</th><th>Protein abundance</th><th>Identified peptide</th><th>Theo. MH+ [Da]</th><th>Residues</th><th>Modifications</th></tr>
<tr><td>Ca-nsLTP1</td><td>A0A2G3AMC5</td><td>Non-specific lipid-transfer protein</td><td>12.0</td><td>9.5</td><td>7.6</td><td>51</td><td>2.28E+08</td><td>TLSGLAQSTDER</td><td>1277.63</td><td>62–73</td><td>None</td></tr>
<tr><td colspan="8" rowspan="3">Sequence:<br>MKAVAVVLVLVVLAAVQLAMA*GRLEQQTGITCGQVDANLAPCVSFLTQGGEPSAACCSGVKTLSGLAQSTDERRTACNCL<br>KAAANRYANLKDDAAQALPGKCGVALNVPISRTINCDT</td><td>TACNCLKAAANR</td><td>1349.64</td><td>75–86</td><td>Carbamidomethyl (C3, C5)</td></tr>
<tr><td>YANLKDDAAQALPGK</td><td>1574.82</td><td>87–101</td><td>None</td></tr>
<tr><td>CGVALNVPISR</td><td>1185.64</td><td>102–112</td><td>Carbamidomethyl (C1)</td></tr>
<tr><td>Ca-nsLTP2</td><td>A0A2G3AAS4</td><td>Non-specific lipid-transfer protein 2</td><td>9.9</td><td>7.1</td><td>8.7</td><td>47</td><td>1.49E+08</td><td>SGSAPSQDCCAR</td><td>1295.51</td><td>43–54</td><td>Carbamidomethyl (C9, C10)</td></tr>
<tr><td colspan="8" rowspan="2">Sequence:<br>MTKGCSFLAVMLVMVLVCNQMGASE*AVTCNPSQLSPCLGALRSGSAPSQDCCARLKGQQSCLCGYMKDPNMKQYVNSP<br>NARKVVGQCGVSVPNC</td><td>GQQSCLCGYMK</td><td>1347.55</td><td>57–67</td><td>Carbamidomethyl (C5, C7)<br>Oxidation (M10)</td></tr>
<tr><td>QYVNSPNAR</td><td>1048.52</td><td>73–81</td><td>None</td></tr>
</table>

For protein identification, the resulting LC–MS/MS spectra were searched using Mascot against the predicted proteome of *Capsicum annuum* cv. proteome. The table reports protein names and accession numbers, theoretical MWs before and after SP removal, mature-protein pIs and sequence coverage, protein abundance estimates, the sequences of the identified tryptic peptides and their theoretical singly protonated peptide masses ($MH^+$). The table also reports the corresponding residue positions and included modifications. In the protein sequences, asterisks indicate the predicted SP cleavage sites and identified peptides are highlighted in red. N-terminal signal peptides (SPs) and their cleavage sites were predicted using SignalP 5.0 (https://services.healthtech.dtu.dk/services/SignalP-5.0/) [2]. Theoretical molecular weights (MWs) and isoelectric points (pI) of the full-length and mature proteins were calculated using ExPASy ProtParam (https://web.expasy.org/protparam/).
